# Parallel Bimodal Single-cell Sequencing of Transcriptome and Chromatin Accessibility

**DOI:** 10.1101/829960

**Authors:** Qiao Rui Xing, Chadi EL Farran, Yao Yi, Tushar Warrier, Pradeep Gautam, James J. Collins, Jian Xu, Hu Li, Li-Feng Zhang, Yuin-Han Loh

**Affiliations:** Epigenetics and Cell Fates Laboratory, Programme in Stem Cell, Regenerative Medicine and Aging, A*STAR Institute of Molecular and Cell Biology, Singapore 138673, Singapore; School of Biological Sciences, Nanyang Technological University, 637551, Singapore; Department of Biological Sciences, National University of Singapore, Singapore 117543, Singapore; Howard Hughes Medical Institute, Boston, MA 02114, USA; Institute for Medical Engineering and Science Department of Biological Engineering, and Synthetic Biology Center, Massachusetts Institute of Technology, Cambridge, MA 02114, USA; Broad Institute of MIT and Harvard, Cambridge, MA 02139, USA; Wyss Institute for Biologically Inspired Engineering, Harvard University, Boston, MA, USA; Center for Individualized Medicine, Department of Molecular Pharmacology & Experimental Therapeutics, Mayo Clinic, Rochester, Minnesota 55905, USA; NUS Graduate School for Integrative Sciences and Engineering, National University of Singapore, 28 Medical Drive, Singapore 117456, Singapore

**Keywords:** Bimodal, ASTAR-Seq, scRNA-seq, scATAC-seq, Transcriptome, Chromatin Accessibility, Co-accessibility

## Abstract

We developed ASTAR-Seq (Assay for Single-cell Transcriptome and Accessibility Regions) integrated with automated microfluidic chips, which allows for parallel sequencing of transcriptome and chromatin accessibility within the same single-cell. Using ASTAR-Seq, we profiled 192 mESCs cultured in serum+LIF and 2i medium, 424 human cell lines including BJ, K562, JK1, and Jurkat, and 480 primary cells undergoing erythroblast differentiation. Integrative analysis using Coupled NMF identified the distinct sub-populations and uncovered sets of regulatory regions and the respective target genes determining their distinctions. Analysis of epigenetic regulomes further unravelled the key transcription factors responsible for the heterogeneity observed.

## INTRODUCTION

With the growing interest in understanding the cellular heterogeneity, development of single cell technologies has exploded in recent years. There are many techniques measuring genome^1^, transcriptome^2^, chromatin accessibility^3^, DNA methylation^4^, chromatin conformation^5^, copy number variation^6^, lineage^7^, and cell surface protein^8^, at a single-cell resolution. scRNA-seq yields comprehensive information of the transcriptome within an individual cell. scATAC-seq enables prediction of the novel cis- and trans-regulatory elements and regulatory transcription factors, providing insights into the regulome heterogeneity^3^. Besides, there are multi-modal methods developed for the concurrent measurement of genome and transcriptome^9,10^, transcriptome and DNA methylome within a single-cell^11,12^. A recent study describes scNMT-seq^13^, which simultaneously measures DNA methylation, chromatin accessibility and gene expression within a single-cell. However, due to the bisulfite treatment, chromatin accessibility libraries suffer from the extremely low mapping percentage, and mutations would be acquired during the treatment. Additionally, scNMT-seq determines accessibility based on the methylation level rather than actual enrichment, rendering it less applicable for the integrative analysis using the pipeline like Coupled NMF, and the prediction of regulatory elements using the software such as CICERO^14^. More recently, two similar bimodal techniques, sci-CAR^15^ and scCAT-seq^16^, are developed to jointly measure chromatin accessibility and gene expression within a single-cell. The former is a combinational indexing-based assay, allowing for profiling of thousands of single cells, whereas the latter is a 96-well PCR plate-based technique. Owing to the high-throughput profiling, sci-CAR libraries suffer from rare sequencing reads, resulting in the extensive signal loss. On the other hand, requisite for the isolation of gDNA and RNA in the first step, makes scCAT-Seq not compatible with the automated platforms, and the extensive manual handling steps render scCAT-Seq not applicable for high-throughput study. Here, we describe an automated Assay for Single-cell Transcriptome and Accessible Regions sequencing (ASTAR-Seq) for concurrent measurement of whole transcriptome and epigenome accessibility of a single-cell with high sensitivity. Multilayers of information collected by ASTAR-Seq allows for the identification of regulatory regions and the genes being regulated, which together contribute to the cellular heterogeneity.

## RESULTS

In ASTAR-Seq, single cells are first captured at different cell capture sites of the Fluidigm C1 microfluidic chip, linked with separate reaction chambers (Fig. 1a). Open chromatins of each cell are then tagmented with Tn5 transposase, from which accessible DNA fragments (ATAC-DNA) with adaptor sequences are generated. Next, mRNA is reverse transcribed to double-stranded cDNA, which is then labeled with biotin during the PCR amplification process. Biotinylation of cDNA enables the separation of cDNA from the ATAC-DNA using the streptavidin beads. Lastly, the separated ATAC-DNA and cDNA fractions are further processed for library preparation and sequenced in parallel. The earlier prototype, where the reverse transcription was performed before the transposition, was not successful (Supplementary Fig. 1a). This could be attributed to Tn5 transposase unexpectedly digesting single-stranded cDNA, which resulted in cDNA to be inseparable from the ATAC-DNA (Supplementary Fig. 1b-d). The current ASTAR-Seq protocol was first optimized and tested with 1000 cells on benchtop. The optimal condition yielded abundant cDNA and achieved clear separation of the ATAC-DNA and cDNA (Supplementary Fig. 1e-g).

**Figure 1.**
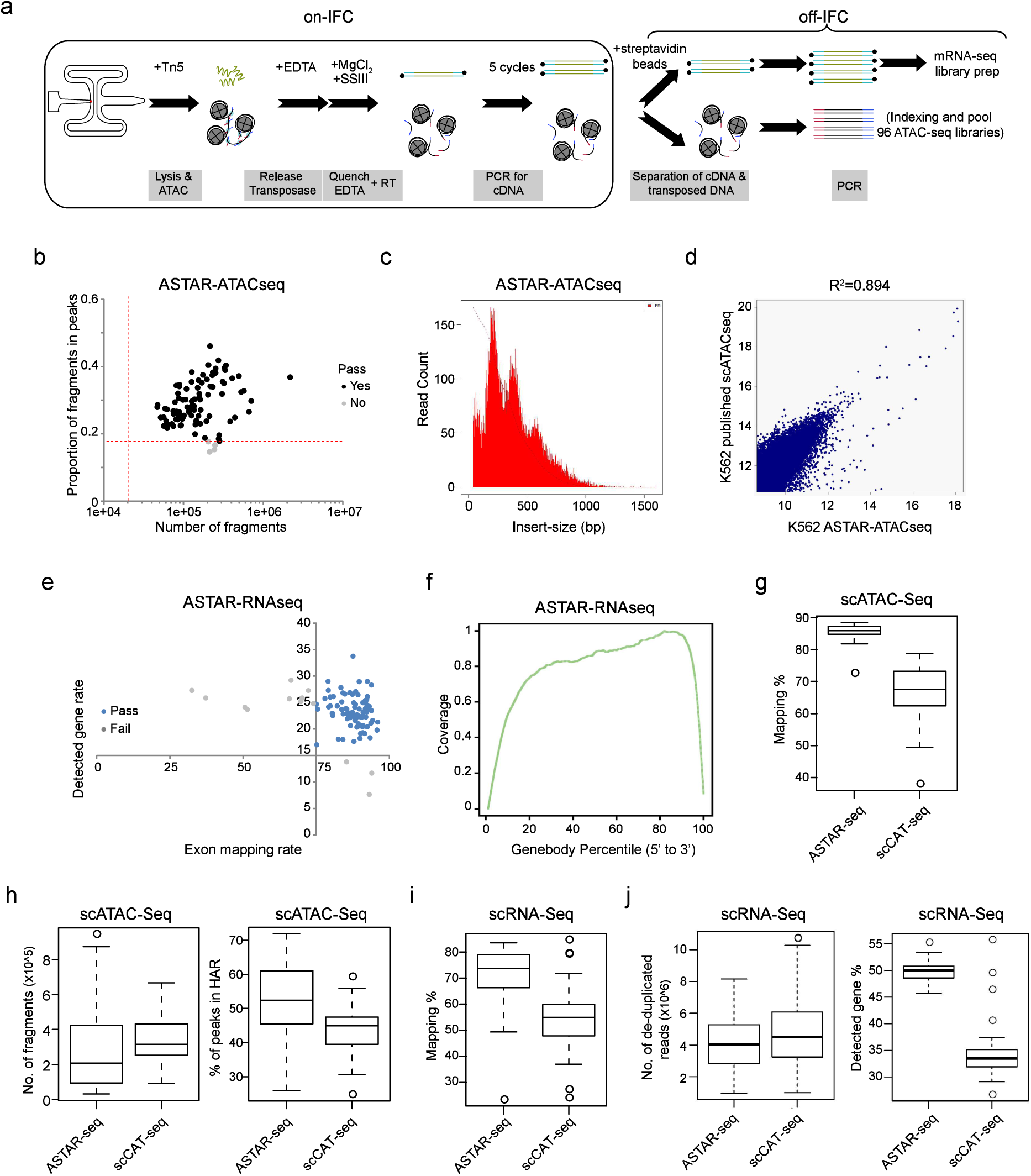
Assay for Single-cell Transcriptome and Accessible chromatin Region (ASTAR-Seq). (a) Overview of the ASTAR**-**Seq protocol. (b) Dotplot revealing the proportion of fragments in peaks (y-axis) against the fragment numbers of each K562 ASTAR-ATACseq library (x-axis). Red dotted lines represent the threshold values set for each criterion. Source data is provided as a Source Data file. (c) Histogram demonstrating the frequency (y-axis) of fragments with the indicated insert size (x-axis). (d) Dotplot demonstrating Pearson correlation between the K562 ASTAR-ATACseq and the published K562 scATACseq^3^ libraries. Source data is provided as a Source Data file. (e) Dotplot revealing the detected gene rate (%) of each K562 ASTAR-RNAseq (y-axis) plotted against its exon mapping rate (x-axis). Blue dots represent the libraries which pass the QC, whereas grey dots represent the libraries of low quality. Source data is provided as a Source Data file. (f) Line plot representing the coverage ratio (y-axis) of K562 ASTAR-RNAseq reads over the genebodies of housekeeping genes (x-axis). (g-h) Boxplots showing the mapping percentage to human genome (g), number of fragments (h-left), and percentage of peaks in HAR (h-right) of scATAC-Seq libraries prepared by scCAT-Seq and ASTAR-Seq protocol. Source data is provided as a Source Data file. (i-j) Boxplots showing the mapping percentage to human genome (i), number of de-duplicated reads (j-left), and detected gene rate (j-right) of scRNA-Seq libraries prepared by scCAT-Seq and ASTAR-Seq protocol. Source data is provided as a Source Data file.

As a proof of concept, we first applied ASTAR-Seq to an ENCODE cell line, K562. Of the 96 ASTAR-ATACseq libraries sequenced, 92 libraries (95.8%) passed the QC thresholds for chromVAR (Fig. 1b and Supplementary Table 1). Of note, a median of 28464 peaks were called, and 27.9% of fragments were in peaks, indicating libraries are of high signal-to-noise profile (Fig. 1b). Insert-size distribution of ASTAR-ATACseq libraries demonstrated clear nucleosomal periodicity, a characteristic pattern of an ATAC-seq library^3^ (Fig. 1c). In addition, ASTAR-ATACseq libraries showed Pearson correlation of 0.89 with the published scATAC-seq libraries^3^, indicative of its high similarity with the unimodal libraries (Fig. 1d). On the other hand, 83 out of 96 ASTAR-RNAseq libraries (86.5%) passed the QC thresholds for detected gene rate (>=15%) and exon mapping rate (>=75%) (Fig. 1e and Supplementary Table 1). scRNA-Seq reads spread across the entire gene body, without biasing towards any end of the mRNA (Fig. 1f).

To make a fair comparison with the similar bimodal technique scCAT-Seq, we sequenced additional 40 K562 ASTAR-Seq libraries at a similar sequencing depth. Specifically, ATAC-Seq libraries prepared by scCAT-Seq protocol displayed significantly lower mapping percentage to the human genome (scCAT-Seq: 67.6%; ASTAR-Seq: 85.8%) (Fig. 1g). Although lesser number of fragments being recovered (scCAT-Seq: 315835; ASTAR-Seq: 213141), ASTAR-Seq outperformed in terms of the percentage of fragments contributing to the highly accessible regions (HARs) (scCAT-Seq: 44.9%; ASTAR-Seq: 52.4%), indicating its better library complexity and higher signal-to-noise ratio (Fig. 1h). Likewise, scRNA-Seq prepared by scCAT-Seq protocol exhibited lower mapping percentage (scCAT-Seq: 54.9%; ASTAR-Seq: 73.8%) (Fig. 1i). Owing to the comparable sequencing depth, similar numbers of de-duplicated reads were detected in both scRNA-Seq libraries (scCAT-Seq: 4507504; ASTAR-Seq: 4047857) (Fig. 1j). Remarkably, ASTAR-Seq exhibited superior performance in terms of the gene detection rate (scCAT-Seq: 33.49%; ASTAR-Seq: 49.94%), suggesting its outstanding sensitivity (Fig. 1j). Taken together, these data indicate the reliability and superior performance of the ASTAR-Seq.

We next applied ASTAR-Seq to 192 E14 mESCs cultured in serum+LIF and 2i medium, which were named as mESCs and 2i cells throughout the study. All the sequenced scATAC-Seq libraries passed the QC thresholds (Fig. 2a and Supplementary Table 2). A median of 87781 peaks were called, and 25% of the fragments contributed to HARs (Fig. 2a and Supplementary Fig. 2a). scATAC-Seq reads were highly enriched at Transcription Start Sites (TSS) and displayed an insert-size distribution with nucleosomal pattern, indications of high quality libraries^3^ (Supplementary Fig. 2b-c). We then clustered the mESCs and 2i ASTAR-ATACseq libraries based on the enrichment of Mouse JASPAR motifs. Although mESCs and 2i cells were mostly clustered separately, a certain degree of overlapping was observed (Fig. 2b). Clustering accuracy was measured using the confusion matrix, which achieved an accuracy of 100% (Supplementary Fig. 2d). Of note, chromatins containing motif sequences of Klf4, Rarg, Zfx, Klf12, and Mlxip showed significant variability in terms of accessibility (P-value <0.05) (Supplementary Fig. 2e and Supplementary Table 2). Klf4 motif was highly accessible in 2i cells, whereas Zfx showed the opposite trend (Supplementary Fig. 2f-g). On the other hand, majority of the ASTAR-RNAseq libraries (80.7%) displayed high gene detection rate and exon mapping rate and demonstrated full gene body coverage for the detected transcripts (Fig. 2c, Supplementary Fig. 2h and Supplementary Table 2). Similarly, confusion matrix also illustrated the highly accurate clustering (95.9%) of mESCs and 2i ASTAR-RNAseq libraries (Supplementary Fig. 2i). To study the heterogeneity within the mESCs, we correlated the ASTAR-RNAseq libraries to the Mouse Cell Atlas (MCA) panel^17^. The analysis revealed three types of cells, including mESCs, ICM-like mESCs, and 2C-like (2-cell) mESCs, in agreement with the previous report^18^ (Fig. 2d). PCA also revealed the separate cluster of minority 2C-like cells (Supplementary Fig. 2j). To confirm the presence of 2C-like cells in our mESCs culture, we utilized the previously constructed *2C::tdTomato* reporter^18^. Indeed, around 1-2% of cells consistently exhibited the activation of the 2C reporter, which was significantly increased upon the depletion of *G9a*, a H3K9me3 methyltransferase, as previously described^18^ (Supplementary Fig. 2k). Intriguingly, MCA analysis of 2i cells exhibited complete absence of 2C-like population (Supplementary Fig. 2l).

**Figure 2.**
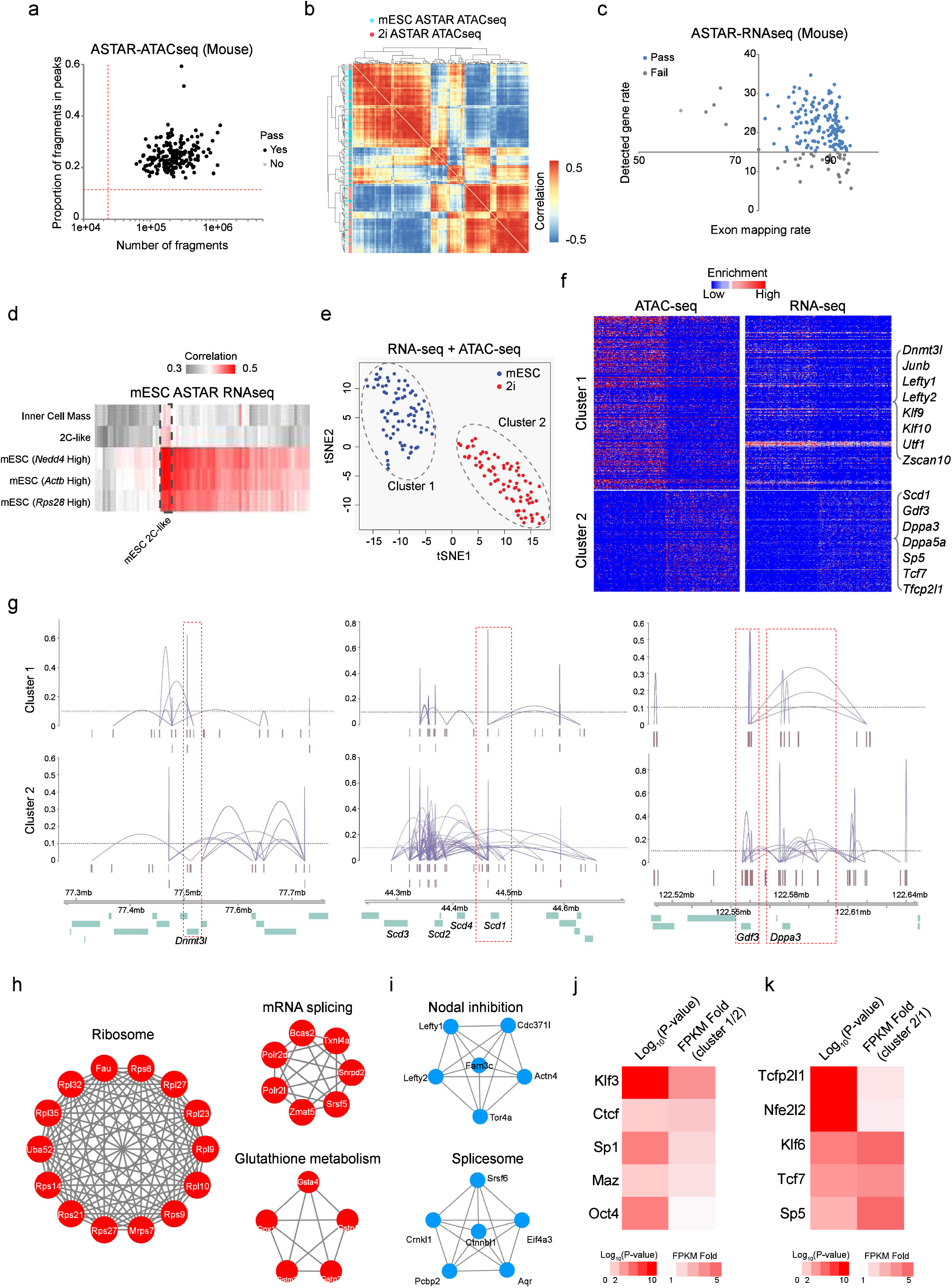
Transcriptomic and Epigenetic Heterogeneity of Primed and Naïve E14 mESCs. (a) Dotplot revealing the proportion of fragments in HARs (y-axis) and number of fragments (x-axis) of each mouse ASTAR-ATACseq library. Red dotted lines represent the thresholds set for each criterion. Source data is provided as a Source Data file. (b) Heatmap demonstrating the correlation among mESCs and 2i cells based on their ASTAR-ATACseq libraries. (c) Dotplot revealing the detected gene rate (%) of each mouse ASTAR-RNAseq (y-axis) plotted against its exon mapping rate (x-axis). Blue dots represent the libraries which pass the QC, whereas grey dots indicate the low-quality libraries. Source data is provided as a Source Data file. (d) Heatmap revealing the correlation of each mESCs ASTAR-RNAseq library to the various lineages of MCA. Color indicates the correlation level, ranging from dark grey (low) to dark red (high). 2-cell like (2C-like) mESCs are boxed with dotted line. Source data is provided as a Source Data file. (e) NMF clustering of mESCs and 2i cells based on the correlative signals of their ASTAR-ATACseq and ASTAR-RNAseq libraries. (f) Heatmaps revealing pairs of accessible regulatory regions (left) and the corresponding genes (right) which are differentially enriched between the NMF clusters. Each column represents an ASTAR-Seq library, whereas each row represents a chromatin region (left) or a gene (right). Color indicates the accessibility (left) and expression (right) levels, ranging from blue (low) to red (high). Representative genes are indicated on the right. Source data is provided as a Source Data file. (g) Line plots demonstrating the differential co-accessibility links between the highlighted regions and its surrounding regions, identified using Cicero. Top plots are constructed from cluster 1 cells, whereas bottom plots are constructed from cluster 2 cells. Peak heights (y-axis) denote the co-accessibility scores. (h-i) Interactome analysis revealing the top pathways enriched with the cluster 1 (i) and cluster 2 (h) specific genes. (j-k) Heatmaps demonstrating the TF motifs enriched with the cluster 1 (j) and cluster 2 (k) specific accessible regions and their relative expressions. Source data is provided as a Source Data file.

We then clustered mESCs and 2i cells using coupled Non-negative Matrix Factorization (NMF)^19^, based on the correlative information of transcriptome and chromatin accessibility of each individual cell. Notably, when clustering was based on either the differentially expressed genes or the accessible regions identified by coupled NMF, two distinct clusters were always observed (Supplementary Fig. 2m). Cluster 1 was mostly comprised of mESCs, whereas cluster 2 was composed of 2i cells. Intriguingly, integrative clustering based on the NMF-specific genes and peaks together demonstrated superior performance in distinguishing the sub-populations, as seen from the clear separation of mESCs and 2i cells (Fig. 2e). Further, the correlation between accessibility and gene expression enabled us to identify regulatory networks specific to each cluster (Fig. 2f and Supplementary Table 2). For example, *Dnmt3l* was highly accessible and expressed in cells of cluster 1, whereas *Scd1, Gdf3*, and *Dppa3* were highly expressed in cluster 2 cells with greater accessibility (Fig. 2f and Supplementary Fig. 2n). In addition, Cicero^14^ co-accessibility analysis demonstrated that *Dnmt3l* gene and its surrounding loci displayed open chromatin architecture with high interaction frequency in cluster 1, on the contrary, more 3D genomic interactions were observed for *Scd1, Gdf3*, and *Dppa3* in cluster 2 (Fig. 2g). This highlights the differential interactive networks where the cluster-specific putative regulatory elements result in the differential expression of the respective genes. Moreover, NMF cluster-specific genes exhibited extensive interaction among themselves, suggesting their involvement in the same biological processes (Supplementary Fig. 2o-p and Supplementary Table 2). Specifically, cluster 2 genes were involved in pathways related to ribosome, mRNA splicing, and glutathione metabolism, which were implicated to play important roles in the acquisition or maintenance of naïve pluripotent state in earlier studies^20,21^ (Fig. 2h and Supplementary Fig. 2p). On the other hand, cluster 1 genes associated with nodal inhibition and spliceosome pathway, which were reported to safeguard the primed pluripotency^22–24^ (Fig. 2i and Supplementary Fig. 2o). To further examine the transcriptions factors responsible for the differential regulatory networks, we performed motif enrichment analysis. Motifs of Klf3, Ctcf, Sp1 and Maz were enriched in cluster 1 specific accessible regions, which were also highly expressed in the cluster 1 cells (Fig. 2j and Supplementary Table 2). Interestingly, an earlier study reported lower genomic looping frequencies in 2i cells as compared to mESCs^25^, supporting the higher Ctcf regulon activity detected in cluster 1. On the contrary, motifs of Tcfp2l1, Nfe2l2, Klf6, Tcf7 and Sp5 were enriched in cluster 2 specific accessible regions, majority of which displayed higher expression in the cells of cluster 2 (Fig. 2k and Supplementary Table 2).

To expand the applicability, we prepared ASTAR-Seq libraries for additional human cell lines, including the BJ cells in adherent culture, and JK1 and Jurkat cells in suspension culture (Figure 3). To characterize the molecular distinction among the hematopoietic cells, 96 K562 libraries sequenced with similar depth were also included for the following analysis. Out of 384 libraries profiled, 375 ASTAR-ATACseq libraries passed the QC thresholds of chromVAR (Supplementary Fig. 3a and Supplementary Table 3). In median, 55193 peaks were called, and 35% fragments contributed to HARs (Supplementary Fig. 3a-b). Insert-size distribution of ASTAR-ATACseq libraries also exhibited characteristic nucleosomal pattern of an ATAC-seq library^3^ (Supplementary Fig. 3c). These indicate the prepared ASTAR-ATACseq libraires are of good quality. In addition, ASTAR-ATACseq libraries also showed a high similarity to the published scATAC-Seq libraries^3^ (Pearson correlation: 0.8). A clustering accuracy of 96.5% was detected using confusion matrix (Supplementary Fig. 3e). Next, we clustered the ASTAR-ATACseq libraries based on the enrichment of human JASPAR motifs, and observed four sub-clusters, among which BJ cells clustered distinctly from the cells of blood lineage and displayed the most distinctive epigenome regulatory profiles (Fig. 3a). Variability analysis identified the transcriptions factors (TFs) defining the cell type identities and their distinctions (Fig. 3b and Supplementary Table 3). Notably, consistent with its distinct cluster, 16 TFs were specifically enriched in BJ cells, such as FOS-JUN and NFE2 (Fig. 3b-d. and Supplementary Fig. 3f). On the other hand, BJ cells also extensively shared motifs with the myeloid lineage cells K562 (79 motifs) and JK1(28 motifs), including ETS1 and ZBTB33 (K562), and families of MEF2, SP, and NFY factors (JK1), as compared to the 6 motifs shared with lymphoid Jurkat cells, such as TEAD family (Fig. 3b-d and Supplementary Fig. 3f). In addition, 46 motifs were shared between the myeloid lineage JK1 and K562 cells, including TFs of GATA-TAL family (Fig. 3b-d).

**Figure 3.**
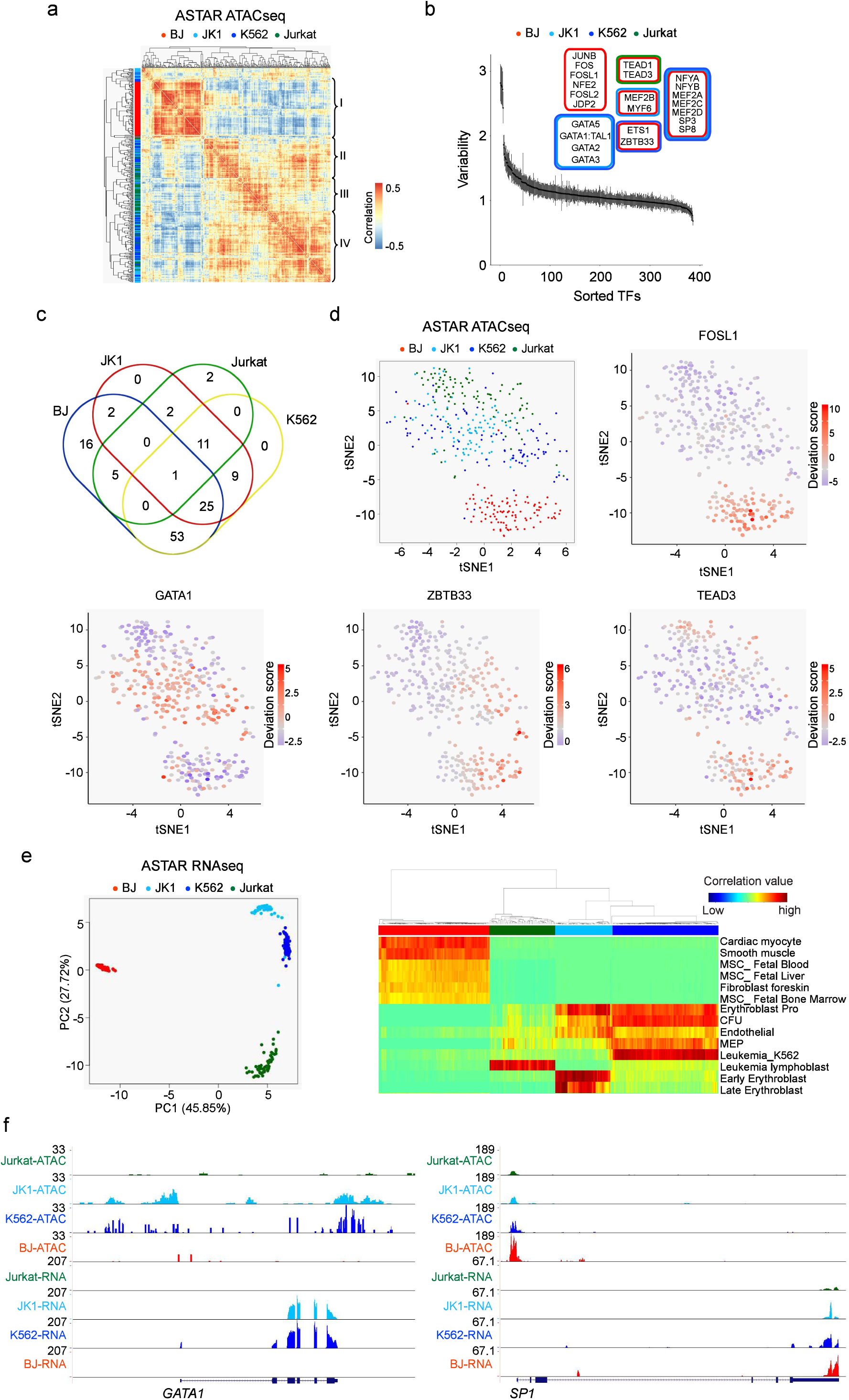
Application of ASTAR-Seq on Human Cell Lines. (a) Clustering of BJ, JK1, K562 and Jurkat ASTAR-ATACseq libraries based on the human JASPAR motif deviation scores calculated over the HARs. Color indicates the correlation level between the libraries, ranging from blue (no) to red (high). Side color bar (y-axis) indicates the identity of each ATACseq library. (b) Variability plot indicating the TF motifs enriched variably across the ASTAR-ATACseq libraries of four human cell lines. Y-axis represents the variability score assigned to each JASPAR motif, whereas x-axis represents the motif rank. Top variable motifs are classified based on their enrichment score across the four cell lines. Box colors indicate the cell lines with high enrichment of TF motifs. Source data is provided as a Source Data file. (c) Multi-Venn diagram showing the shared and unique TFs across the cell lines. (d) Top left: tSNE clustering of BJ, JK1, K562 and Jurkat ASTAR-ATACseq libraries based on the deviation scores of human JASPAR motifs. Colors represent the time-points. Top right & bottom: super-imposition of the motif enrichment scores for FOSL1, GATA1, ZBTB33, and TEAD3 on the tSNE cluster. Colors represent the motif enrichment level, ranging from blue (no) to red (high). (e) Left: PCA clustering of ASTAR-RNAseq libraries based on their correlation to the RCA panel. Right: Heatmap showing the lineages that each cell correlate to. Source data is provided as a Source Data file. (f) UCSC screenshots indicating the chromatin accessibility levels (top panels) and the expression (bottom panels) of *GATA1* (left) and *SP1* (right) across the cell lines.

Of the 384 ASTAR-RNAseq libraries, 296 libraries (77.1%) passed the QC and exhibited full coverage for the expressed transcripts (Supplementary Fig. 3g-h and Supplementary Table 3). Of the 296 cells with RNA-seq library of good quality, only 5 cells failed the scATAC-seq QC (1.69%) (Supplementary Table 3). In summary, a total of 291 cells (75.8%) passed the QC filtration for both scATAC-seq and scRNA-seq (Supplementary Table 3). Meta-analysis of K562 ASTAR-RNAseq, K562 bulk RNA-seq, BJ ASTAR-RNAseq and BJ scRNA-seq illustrated the expected clustering according to the respective cell types^26^ and indicated the similarity of human ASTAR-RNAseq libraries to the bulk RNA-seq or unimodal scRNA-seq previously generated (Supplementary Fig. 3i). Clustering accuracy of the ASTAR-RNAseq libraries was again measured by confusion matrix, which reached an accuracy of 99.63% (Supplementary Fig. 3j). Expectedly, Reference Component Analysis (RCA) analysis demonstrated distinct correlation of BJ cells to the foreskin fibroblasts and muscle lineage cells, and Jurkat cells to the leukemia lymphoblasts (Fig. 3e). Of note, K562 cells showed highest correlation to leukemia K562 and erythroblast progenitors, whereas JK1 exhibited correlation to the erythroblast progenitors, early and late erythroblast (Fig. 3e). Consistent with their differential regulatory activities, GATA1 was only accessible and expressed in JK1 and K562 cells, whereas SP1 was accessible and expressed in BJ, JK1, and K562 cells (Fig. 3f). Taken together, ASTAR-RNAseq libraries of the human cell lines are of good quality, differential analysis of which identified the TFs responsible for the cell type distinctions.

To measure its capability in capturing the dynamic changes, we applied ASTAR-Seq to the primary cells undergoing erythroblast differentiation. We harvested a total of 480 cells at day 6, day 8, day 10, and day12 of erythroblast differentiation, which was induced from the mononuclear cells isolated from umbilical cord blood, and subjected them to ASTAR-Seq library preparation (Fig. 4a). Of the 480 cells, 273 cells (56.9%) demonstrated good quality libraries for both scATAC-seq and scRNA-seq, whereas 41 cells (8.5%), 103 cells (21.5%), and 63 cells (13.1%) failed the QC for scATAC-seq, scRNA-seq, and both respectively (Supplementary Table 4). We next investigated trajectories of the erythroblast differentiation process using pseudotemporal analysis^27,28^. The resultant trajectories consisted of 2 branching events and 5 states (Fig. 4b-c). Interestingly, pseudotime highly correlated with the actual differentiation time points. For instance, cells of early time points such as D6 and D8 were mostly at state 1-3, and D10 cells were mostly at state 4 and 5, whereas majority of D12 cells belonged to state 5 and located at the end of pseudotime (Fig. 4b-c). Expectedly, *HBA2*, a hemoglobin gene, showed elevated expression in cells of state 5 as compared to the others. Next, RCA analysis was performed to track the differentiation status of cells of various states. Notably, majority of the cells exhibited strong correlation to the myeloid or erythroid lineages (Fig. 4d). As the time-point increased, transitions from common myeloid progenitors (CMPs) to myeloid erythroid progenitors (MEPs), and from erythroblast progenitor to early erythroblast and to late erythroblast were observed. Specifically, majority of state 1-3 cells showed strong correlation to the erythroblast progenitor cells and some degrees of correlation to the MEPs and early erythroid cells (Fig. 4d). Cells of state 5 can be broadly classified into three groups, including groups with correlation to early erythroid cells but not to MEPs, with strong early erythroid identity, and with significant late erythroid fate (Fig. 4d). On the other hand, cells of state 4 displayed significant correlation to granulocyte monocyte progenitor (GMP). In addition, state 5 cells abundantly expressed genes related to oxygen transport, hydrogen peroxide catabolic process, cell cycle arrest and erythrocyte differentiation process, whereas state 4 cells expressed genes associated with innate immune response, antigen processing and presentation via MHC class II process (Supplementary Fig. 4a). On the contrary, as compared to state 1 cells, genes that were repressed in state 5 cells were involved in stem cell population maintenance, positive regulation of H3-K4 methylation, and chromatin organization, whilst genes related to erythrocyte maturation, oxygen transport and NF-KB signaling were repressed in state 4 cells (Supplementary Fig. 4b). In conclusion, state 5 cells represent the cells attaining erythroid fate, whereas state 4 cells deviate from the differentiation path to erythroblast, instead acquire the GMP identity.

**Figure 4.**
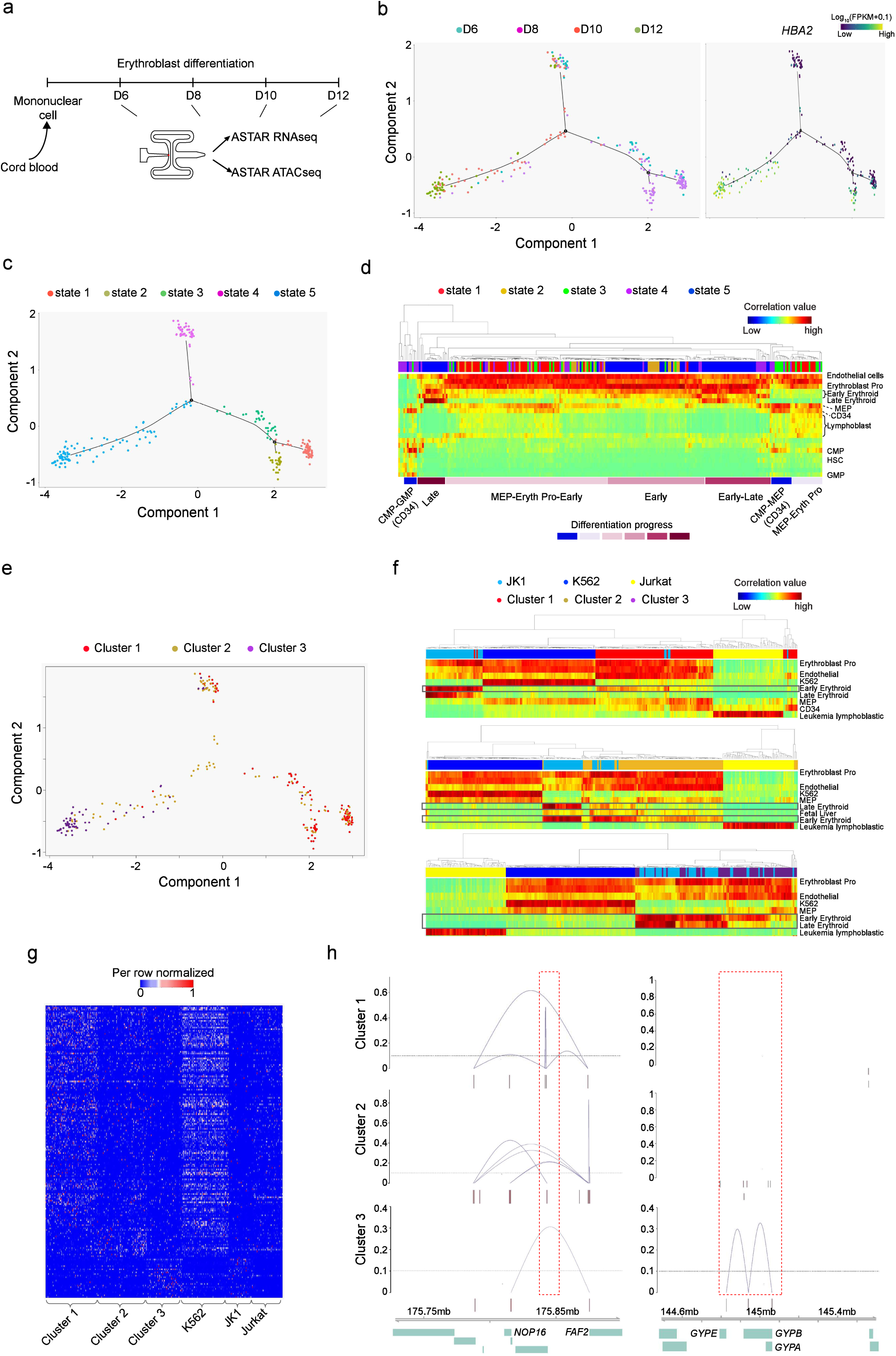
Interactive Analysis of Transcriptomic and Chromatin Accessibility during the Erythroblast Differentiation. (a) Schematic of the erythroblast differentiation time-points harvested for ASTAR**-**Seq library preparation. (b) Left: trajectory of erythroblast differentiation identified by the DDRTree dimension reduction of ASTAR-RNAseq libraries. Right: Superimposition of *HBA2* expression on the trajectory. (c) Trajectory plots indicating the pseudotemporal states. (d) RCA clustering heatmap of the cells undergoing erythroblast differentiation, based on their correlation with the cells of different lineage origins in the RCA panel. Color indicates correlation value, ranging from blue (low) to red (high). Each row indicates one lineage, whereas each column represents an ASTAR-RNAseq library. Pseudotemporal state of each library is indicated on top. Cellular differentiation status is determined based on their correlation to the cells of RCA panel, and indicated below. Source data is provided as a Source Data file. (e) Superimposition of NMF clusters on the trajectory plots of erythroblast differentiation. (f) RCA clustering of JK1, K562, Jurkat cells and the cells of NMF cluster 1 (top), cluster 2 (middle) and cluster 3 (below), respectively. Color indicates correlation value, ranging from blue (low) to red (high). Each row indicates one lineage, whereas each column represents an ASTAR-RNAseq library. Cellular identities are indicated on top. Source data is provided as a Source Data file. (g) Heatmap demonstrating the expression of differential genes across the NMF clusters and the hematopoietic cell lines. Color indicates the expression level, ranging from blue (no) to red (high). Source data is provided as a Source Data file. (h) Line plots indicating the Cicero co-accessibility links between the regions highlighted in red and the distal sites in the surrounding region. The height indicates the Cicero co-accessibility score between the connected peaks. The links are constructed from cells of cluster 1 (top), cluster 2 (middle) and cluster 3 (bottom).

To identify the genes and regulatory regions responsible for the differentiation progression, we applied coupled NMF analysis and identified three clusters. Superimposition of NMF clusters on the erythroblast differentiation trajectory demonstrated that majority of cluster 1 cells belonged to state 1-2, whereas cells of cluster 3 were the major constituent of state 5. Cluster 2 cells scattered across the various states (Fig. 4e). Consistent with the identity of pseudotemporal states, cluster 1 cells mostly exhibited pro-erythroblast and early erythroid characteristics, and clustered closer to K562 cells (Fig. 4f). Cluster 2 cells showed stronger early erythroid identity and started to develop late erythroid characteristics. Notably, cluster 2 cells demonstrated higher similarity to JK1 than K562 cells, possibly reflecting their maturity (Fig. 4f). Indeed, studies suggested that K562 resemble early erythroid precursor cells^29^. On the contrary, cluster 3 cells displayed strong late erythroid identity and were clustered intimately with JK1 cells (Fig. 4f). Supporting this notion, differentially expressed genes of NMF cluster 1 were highly represented in K562 cells, whereas cluster 3 genes were specifically expressed in JK1 cells (Fig. 4g). This was further substantiated using CTen^30^ (Supplementary Fig. 4c-d). On the other hand, chromatins containing motif sequences of GATA1:TAL1 and MEF2D were highly variable in terms of accessibility across the clusters (Supplementary Fig. 4e-f and Supplementary Table 4). In addition, cluster 1 accessible regions were enriched with HOXB4 and FOXA3 motifs, cluster 2 with HNF6 motif, and cluster 3 with OCT2 and RUNX2 motifs, implicating their importance in regulating the genes crucial for the progression of erythroblast differentiation (Supplementary Fig. 4g and Supplementary Table 4). Moreover, the cluster specific genes demonstrated differential co-accessibility with its surrounding regulatory regions. For example, *NOP16* was highly expressed and accessible in cluster 1 with multiple genomic interactions with the nearby genes, whereas cluster 3 specific gene *GYPB* demonstrated high interaction frequency with *GYPE* only in cluster 3 (Fig. 4h).

Altogether, we present an automated bimodal technology ASTAR-Seq, which enables parallel profiling of transcriptome and chromatin accessibility within the same single-cell, at a greater sensitivity. ASTAR-Seq is a powerful integrated approach to understand the connectivity between transcription and epigenetic regulation. We expect the technology to have attractive applications in early embryonic testing, identification of rare cancer sub-populations, and atlasing of whole tissues or cell types.

## METHODS

Methods and any associated references are available in the supplementary information.

## DATA AVAILABILITY

All the raw data has been deposited to GEO database: GSE113418. The source data underlying Figs 1b, 1d, 1e, 1g, 1h, 1i, 1j, 2a, 2c, 2d, 2f, 2j, 2k, 3b, 3e, 4d, 4f, 4g and Supplementary Figs 1b, 1d-g, 2a, 2e, 2l, 3a, 3b, 3d, 3g, 4a, 4b, 4e are provided as Supplementary File 1 under ‘Source Data’ sub-folder. The scripts for bioinformatics analysis were provided as Supplementary File 1 under ‘Bioinformatic scripts’ sub-folder. The ASTAR-Seq scripts of the microfluidic chip (Fluidigm) were provided as Supplementary File 1 under ‘ASTAR script’ sub-folder. All the relevant data are available within the manuscript and Supplementary files or from the corresponding author upon request.

## ACKNOWLEDGMENTS

We are grateful to Haitong Fang, Naresh Waran Gnanasegaran, Yingying Zeng and Sudhagar Samydurai for technical assistance. H.L. is supported by the funding from the National Institutes of [Health CA196631-01A1] and [1U54GM114838-01]. Y-H.L. is supported by the [JCO Development Programme Grant - 1534n00153] and [NRF Investigatorship grant]. Y-H.L. and L-F.Z. are supported by the Singapore National Research Foundation under its Cooperative Basic Research Grant administered by the Singapore Ministry of Health’s National Medical Research Council [NMRC/CBRG/0092/2015]. We are grateful to the Biomedical Research Council, Agency for Science, Technology and Research, Singapore for research funding.

## AUTHOR CONTRIBUTIONS

Contribution: Q.R.X developed the protocol, performed experiments and wrote the paper. C.E.F performed all bioinformatics analyses of the data and wrote the paper; Y.Y, P.G, T.W assisted with method development and prepared text for the paper; J.C, J.X, H.L, L.F.Z analyzed data; and Y.H.L. designed the method, analyzed data, and wrote the paper.

