## Supplemental Information for "Parallel Bimodal Single-cell Sequencing of Transcriptome and Chromatin Accessibility"

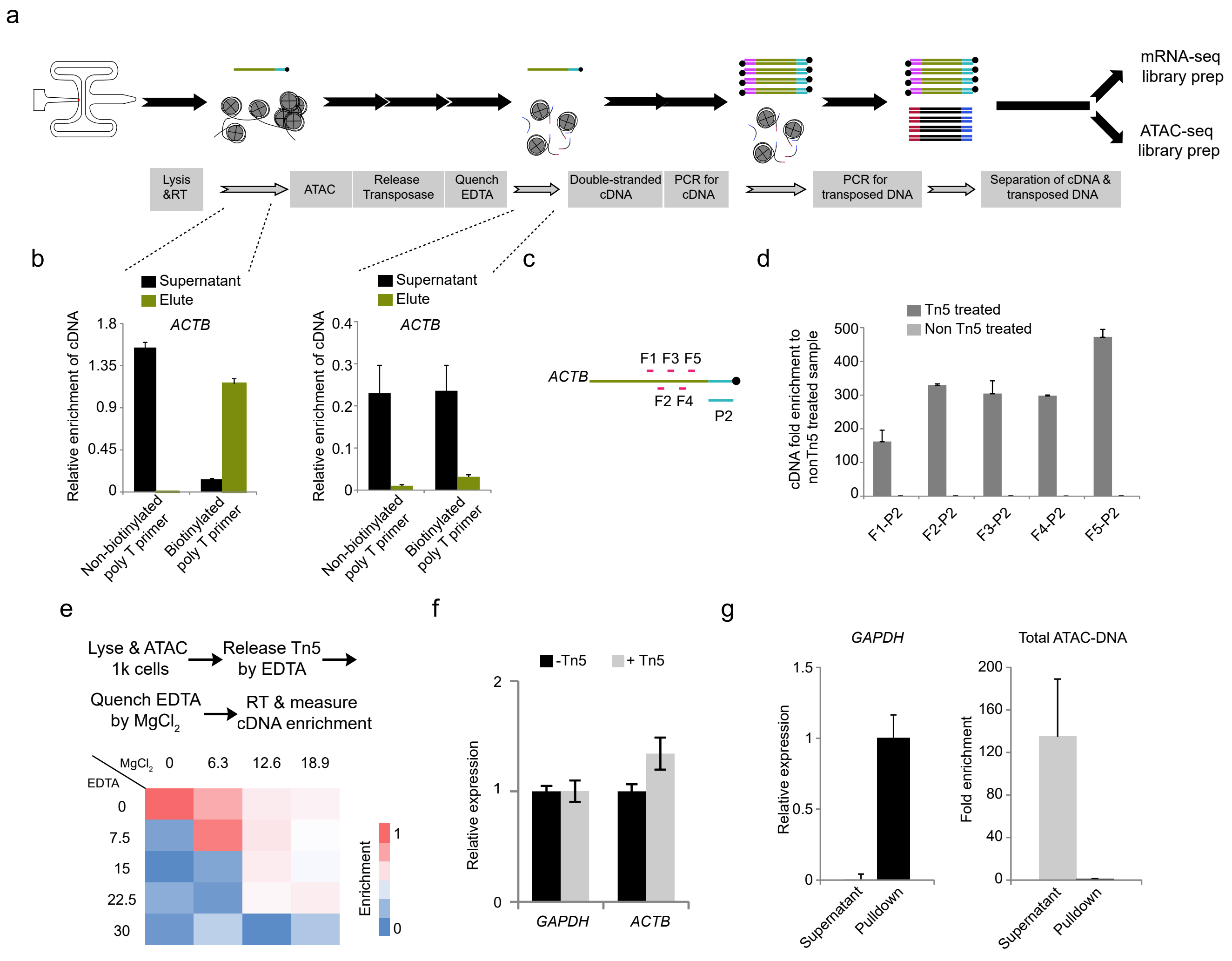


Supplementary Figure 1

**Supplementary Figure 1. Earlier prototype and optimization of ASTAR-Seq platform.**

(a) Overview of the un-optimized ASTAR-seq prototype. Briefly, cells were lysed and mRNA was reverse transcribed to single-stranded cDNA with biotin tag. Open chromatin was tagmented by transposases Tn5 followed by the inactivation of its enzymatic activities. Next, the single-stranded cDNA was converted to double strands and further amplified by biotinylated primers. After which, the tagmented chromatin was amplified by non-biotinylated adaptors which was attached to transposases initially and integrated to open chromatin regions during the transposition reaction. Lastly, open chromatin and biotinylated cDNA were separated by streptavidin beads and proceed for library preparation. (b) Left: Barchart showing the relative enrichment of *ACTB* (Supplementary Table 5) in the supernatant and eluent of the samples right after the reverse transcription. Streptavidin beads pulled down most of the biotinylated cDNA, while non-biotinylated cDNA was mostly in the supernatant, suggesting the specificity of the streptavidin beads. Error bar indicates SD, n=2. Source data is provided as a Source Data file. Right: Barchart showing the relative enrichment of *ACTB* in the supernatant and eluent of the samples right after the inactivation of Tn5. Most of the biotinylated cDNA was not in the eluent, an indication of Tn5 digesting the single-stranded cDNA. Error bar indicates SD, n=2. Source data is provided as a Source Data file. (c) Schematic of primer design on *ACTB* to confirm the digestion of single-stranded cDNA by Tn5. If single-stranded cDNA was digested by Tn5, the digested fragments of cDNA would be further amplified by ATAC adaptors. (d)

Barchart showing the relative enrichment of *ACTB* in the Tn5 treated sample to the non-Tn5 treated control after PCR amplification of the tagmented open chromatin by the ATAC adaptors (Supplementary Table 5). *ACTB* primers F1-F5 (Supplementary Table 5) were used as forward primer, and primer C1-P2-PCR (Supplementary Table 5) with the same sequence as poly T primers with exclusion of poly T sequence, was used as reverse primer. Indeed, single-stranded cDNA was digested by Tn5, as fragments of cDNA could be amplified by the ATAC adaptors. Error bar indicates SD, n=2. Source data are provided as a Source Data file. (e) Heatmap showing the enrichment of *ACTB* after inactivating the Tn5 activity by different dosages of EDTA and quenching excess EDTA by variable amounts of $MgCl_2$. The schematic of the experimental design is showed on top. Source data is provided as a Source Data file. (f) Barchart showing the relative enrichment of *GAPDH* (Supplementary Table 5) and *ACTB* in the samples processed by the pipeline in (e), with or without addition of Tn5. The cDNA amount in the Tn5 treated sample was comparable to the non-treated sample. Error bar indicates SD, n=2. Source data is provided as a Source Data file. (g) Barchart showing the relative enrichment of *GAPDH* (left) and the total amount of ATAC-DNA (right) in the supernatant and eluent of 1000 BJ cells processed by the pipeline in Figure 1a. Error bar indicates SD, n=2. Source data is provided as a Source Data file.

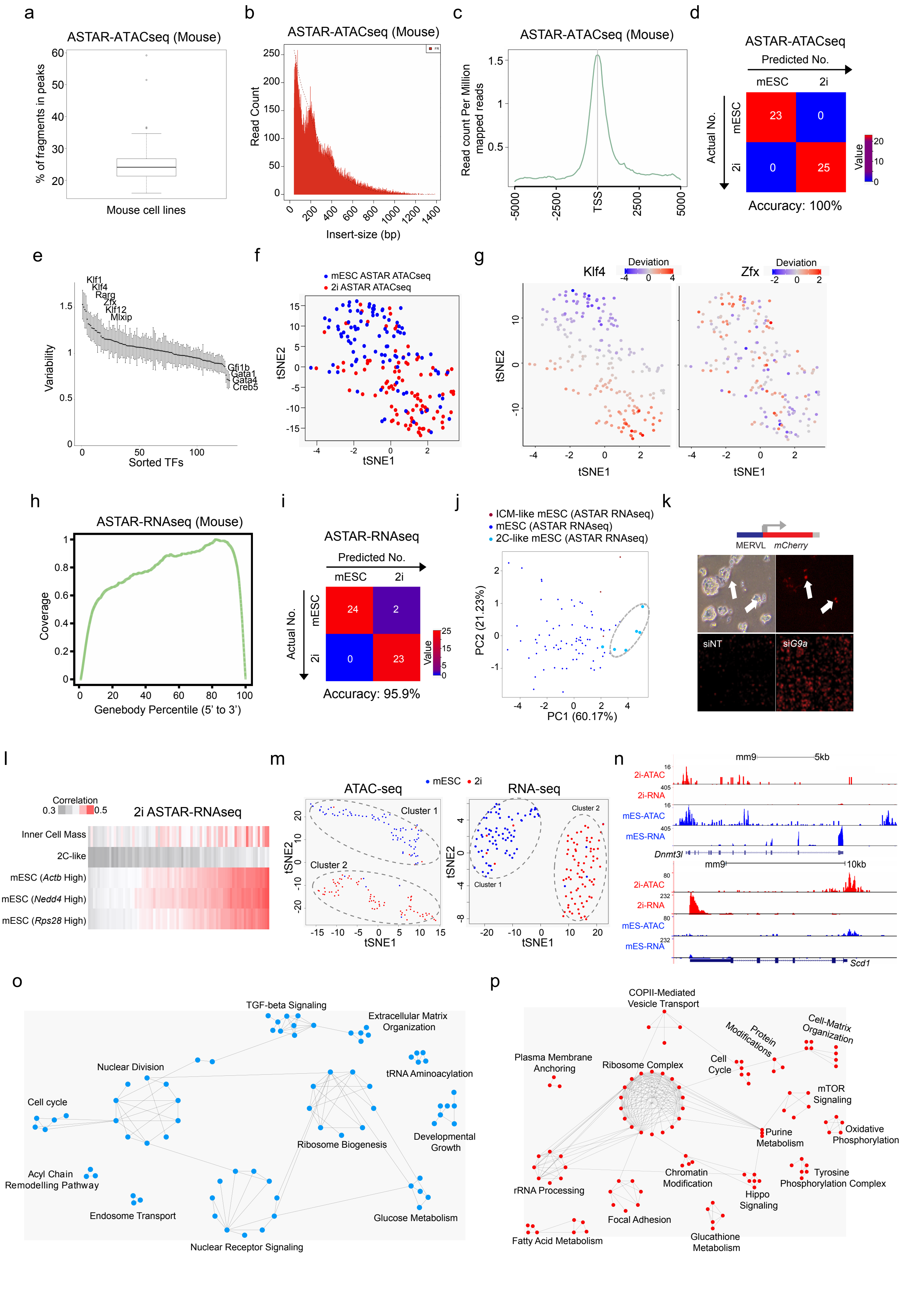


Supplementary Figure 2

**Supplementary Figure 2. Heterogeneity of mESC and 2i cells in terms of chromatin accessibility and transcriptome.**

(a) Boxplot showing the percentage of the fragments of each mouse ASTAR-ATACseq library contributing to HARs (Highly Accessible Regions). Source data is provided as a Source Data file. (b) Histogram demonstrating the frequency (y-axis) of fragments with the indicated insert size (x-axis). (c) Average enrichment profile indicating the read counts per million mapped reads of a mouse ASTAR-ATACseq library, around all transcription start sites (TSS) of the genome with a window of -5K to 5K. (d) Confusion matrix for ASTAR-ATACseq libraries of mESCs and 2i cells using Support Vector Machines with Radial Basis Function Kernel algorithm. (e) Line plot showing the variability level (y-axis) of all mouse JASPAR motifs on the determined HARs of the mESCs and 2i cells. Source data is provided as a Source Data file. (f-g) tSNE clustering of mouse ASTAR-ATACseq libraries (f) and the superimposition of the deviation scores of Klf4 (g-left) and Zfx (g-right) motifs on the tSNE cluster. (h) Line plot representing the coverage ratio (y-axis) of K562 ASTAR-RNAseq reads over the genebodies of housekeeping genes (x-axis). (i) Confusion matrix for ASTAR-RNAseq libraries of mESCs and 2i cells using Random Forest algorithm. (j) PCA clustering of the mESCs ASTAR-RNAseq libraries based on MCA correlation levels. The dotted ellipse surrounds the 2C-like mESCs. (k) Schematic of the 2C reporter (top), and bright field (top left) and fluorescence (top right) images of mESCs transfected with 2C reporter. White arrows indicate the cells in which the 2C reporter is activated. Fluorescence images of siNT control mESCs (bottom left) and *G9a*-depleted mESCs (bottom right) transfected with the 2C reporter is shown. (l) Heatmap revealing the correlation levels of 2i ASTAR-RNAseq libraries with the indicated lineages using MCA. Color indicates the correlation level, ranging from dark grey (low) to dark red (high). Source data is provided as a Source Data file. (m) tSNE clustering of mESCs and 2i ASTAR-ATACseq libraries (left) and ASTAR-RNAseq (right) libraries based on the differential accessible chromatin regions and differentially expressed genes identified by NMF clustering. (n) UCSC screenshots demonstrating the chromatin accessibility and expression levels of the genes that are differentially accessible and expressed between the NMF clusters. (o-p) Interactome analysis showing the strong interaction among the cluster 1 (o) and cluster 2 (p) specific genes, identified by NMF analysis, in the specified pathways.

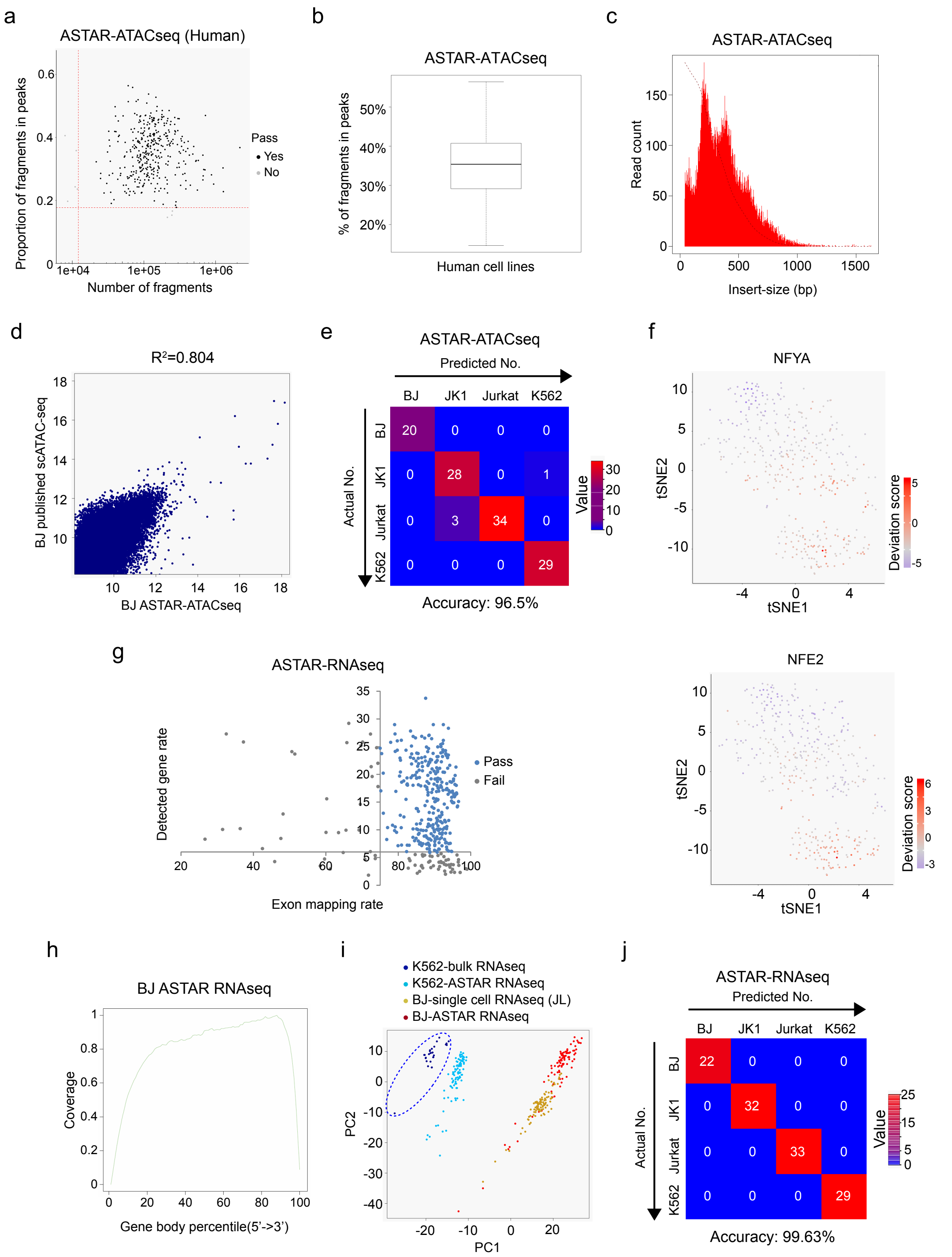


Supplementary Figure 3

**Supplementary Figure 3. Quality control of human ASTAR-Seq libraries.**

(a) Dotplot revealing the proportion of fragments of each human ASTAR-ATACseq library that falls in the HARs (y-axis) plotted against the size of each library (x-axis). Red dotted lines represent the threshold for each criterion. Source data is provided as a Source Data file. (b) Boxplot showing the percentage of fragments of each human ASTAR-ATACseq library contributing to HAR (Highly Accessible Regions). Source data is provided as a Source Data file. (c) Histogram demonstrating the frequency (y-axis) of fragments with the indicated insert size (x-axis). (d) Pearson correlation between BJ ASTAR-ATACseq and published BJ scATACseq[3], $R^2$ is 0.804. The axes represent the accessibility level in each sample. Source data is provided as a Source Data file. (e) Confusion matrix for ASTAR-ATACseq libraries of the human cell lines using Random Forest algorithm. (f) Super-imposition of the motif enrichment scores of NFYA and NFE2 on the tSNE cluster in Figure 3d. Color indicates the enrichment level, ranging from blue (no) to red (high). (g) Dotplot revealing the detected gene rate (%) of each human ASTAR-RNAseq library (y-axis) plotted against the rate of mapping to exons (x-axis). Blue dots represent the libraries which pass the QC, whereas the grey dots represent the libraries that need to be filtered out. Source data is provided as a Source Data file. (h) Line plot representing the coverage ratio (y-axis) of the human ASTAR-RNAseq libraries over the genebodies of housekeeping genes (x-axis). (i) PCA of K562 bulk RNAseq (dark blue, dotted circles), K562 ASTAR-RNAseq (light blue), BJ single cell RNAseq (golden yellow) and BJ ASTAR-RNAseq (red) libraries. BJ and K562 ASTAR-RNAseq libraries display similar transcriptomic profile to conventional BJ and K562 RNAseq libraries respectively. (j) Confusion matrix for ASTAR-RNAseq libraries of the human cell lines using Random Forest algorithm.

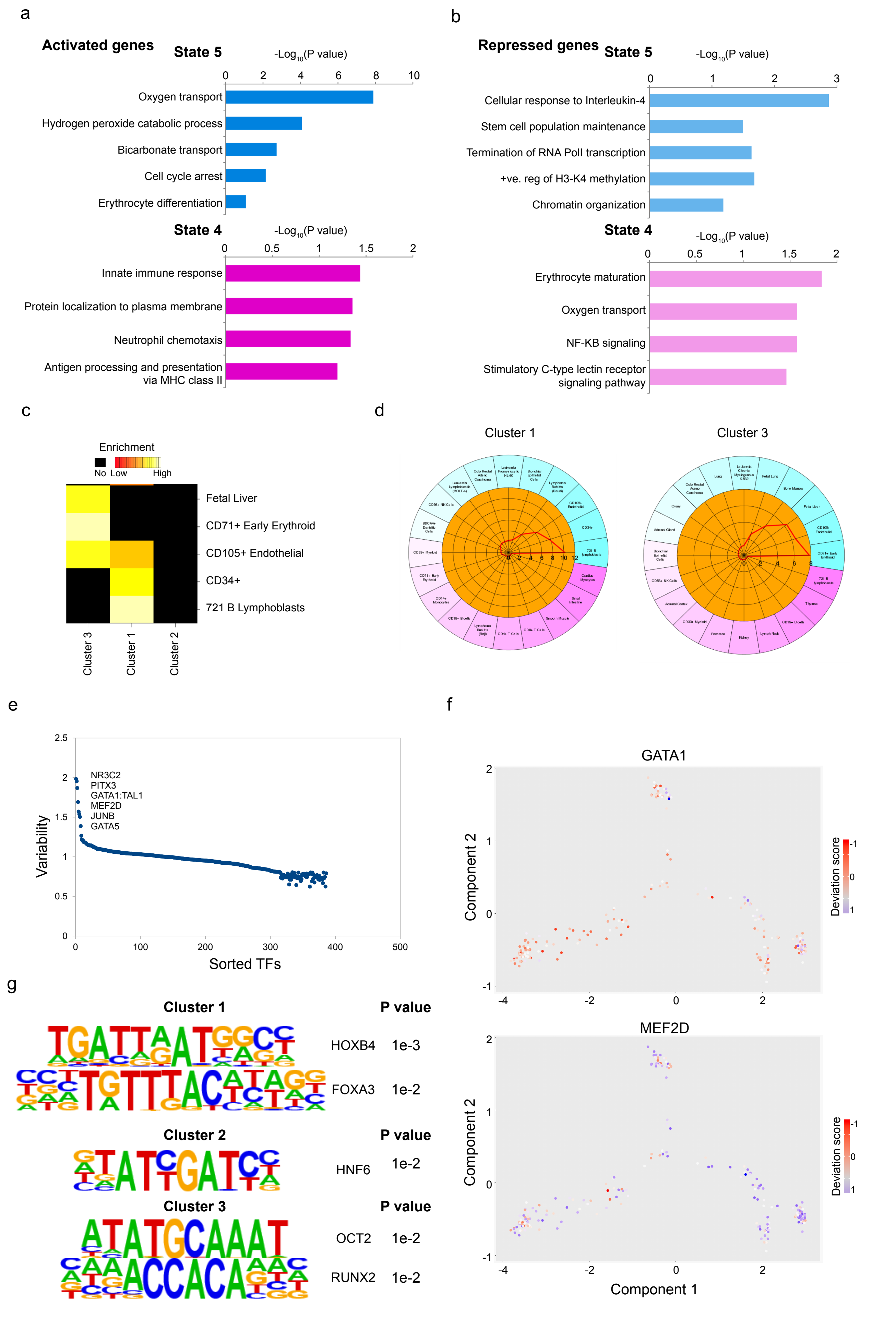


Supplementary Figure 4

**Supplementary Figure 4. Characterization of cells of each NMF cluster.**

(a-b) Barchart showing the Gene Ontology terms enriched by the activated (a) or repressed (b) genes in cells of state 5 (top) or state 4 (bottom) as compared to the cells of state 1. Source data is provided as a Source Data file. (c) Heatmap demonstrating the cell lineages enriched with the NMF cluster specific genes, analyzed by CTEN. Color represents the enrichment score, ranging from black (no) to red (low) and yellow (high). (d) Pie charts showing the lineages enriched with the NMF cluster 1 (left) and cluster 3 (right) genes. (e) Dot plot showing the variability level (y-axis) of all human JASPAR motifs on the determined HARs of the cells undergoing erythroblast differentiation. Source data is provided as a Source Data file. (f) Super-imposition of the motif enrichment scores for GATA1 and MEF2D on the trajectory of erythroblast differentiation. Color represents the enrichment level, ranging from blue (no) to red (high). (g) Transcription factor motifs enriched with the identified cluster 1 (top), cluster 2 (middle) and cluster 3 (bottom) specific accessible regions. P values are indicated on the right.

### SUPPLEMENTARY EXPERIMENTAL PROCEDURES

#### Cell lines culture

mES-E14TG2a mouse embryonic stem cells were maintained in DME+4500 mg/l medium (HyClone) supplemented with 15% Fetal Bovine Serum (Gibco), MEM Non-Essential Amino Acids Solution (100X, Gibco), 200mM L-Glutamine (100X, Gibco), 55mM β-mercaptoethanol, 100 U/ml Penicillin-Streptomycin (100X, Gibco), 10^7 unit/ml ESGRO® Recombinant Mouse LIF Protein (10000X, Merck).

2i cells were derived by culturing mES-E14TG2a cells with N2B27 based 2i medium and harvested for ASTAR-seq at passage 3. The recipe of N2B27 based 2i medium is as follows: DMEM/F12 medium (Gibco), Neurobasal medium (Gibco), N2 Supplement (200X, Gibco), B27 Supplement (100X, Gibco), 200mM L-Glutamine (200X, Gibco), 0.1mM β-mercaptoethanol (Gibco), 7.5% BSA (1500X, HyClone), 3uM CHIR99021 (STEMCELL Technologies), 1uM PD0325901 (STEMCELL Technologies), 10^7 unit/ml ESGRO® Recombinant Mouse LIF Protein (10000X, Merck). Cells were routinely propagated by trypsinization and re-plated onto 0.1% gelatin coated plates every three to four days.

Human neonatal fibroblast cell line, BJ (Stemgent, Cambridge, MA), were maintained in BJ medium, which is composed of DME+4500 mg/l medium (HyClone) supplemented with 10% Fetal Bovine Serum (Gibco), MEM Non-Essential Amino Acids Solution (100X, Gibco), 200mM L-Glutamine (100X, Gibco), 10000 U/ml Penicillin-Streptomycin (100X, Gibco). K562 (ATCC® CCL-243™) were maintained in K562 medium, which is composed of RPMI1640 medium with L- Glutamine (HyClone) supplemented with 10% Fetal Bovine Serum (Gibco), 200mM L-Glutamine (100X, Gibco), 10000 U/ml Penicillin-Streptomycin (100X, Gibco). Jurkat, Clone E6-1 (ATCC® TIB-152™) were maintained in RPM1640 medium (Hyclone) supplemented with 10% Fetal Bovine Serum (Gibco). JK-1 (ACC 347) were maintained in RPMI1640 medium (Hyclone) supplemented with 20% Fetal Bovine Serum (Gibco).

#### Isolation of Mononuclear cells and erythroblast differentiation

The umbilical cord blood samples were collected from Singapore Cord Blood Bank (SCBB; NUS IRB B-15-051) and the mononuclear cells were isolated via density gradient centrifugation using Ficoll® Paque Plus as described by the manufacturer

(GE Healthcare, Uppsala, Sweden).

The mononuclear cells were cultured in Serum Free Expansion Medium II (SFEM II, Stem Cell Technologies, Vancouver, Canada) with recombinant human interleukin-3 (10ng/mL; Peprotech, NJ, USA), recombinant human interleukin-6 (10ng/mL; Peprotech, NJ, USA), recombinant human stem cell factor (50ng/mL; Peprotech, NJ, USA), Recombinant human erythropoietin (2U/mL; R&D System, MN, USA), recombinant human insulin-like growth factor 1 (40ng/mL; Peprotech, NJ, USA), Dexamethasone (1mM; Sigma-Aldrich, MO, USA), and L-ascorbic acid (50ug/mL; Sigma-Aldrich, MO, USA). The media was replenished every alternate day for a period of 14 days in 24-well plate culture format.

**Dead cell exclusion**

Dead cells were excluded from the cells undergoing erythroblast differentiation, following the instructions of Dead Cell Removal Kit (Miltenyi Biotec). The live cells were collected for ASTAR-seq library preparation.

**ASTAR-seq METHOD**

**1. on-IFC procedures of ASTAR-seq**

We used the C1 single-cell Auto Prep System with its Open App™ program (Fluidigm) and a novel and cognate ASTAR-seq pipeline to prepare ATACseq and mRNAseq libraries from an individual cell. Single cells were captured on the C1 Single-Cell Auto Prep IFC microfluidic chip using the 'ASTAR: Cell Load (1861x/1862x/1863x)' script, generated using the C1™ Script Builder software. Firstly, cells were trypsinized, washed once with C1 Cell Wash Buffer (Fluidigm), diluted to 300-600 cells/µL, and mixed with C1 Cell Suspension Reagent at a ratio of 3:2. Then, 20 µL of the cell mix was loaded on to the Fluidigm IFC for cell capture. Optionally, the 'ASTAR: Cell Load and stain (1861x/1862x/1863x)' script can be used when LIVE/DEAD viability dyes (1:1000 of green-fluorescent Calcein-AM dye, and 1:1000 red-fluorescent ethidium homodimer-1 dye (Life Technologies) diluted with C1 cell wash buffer (Fluidigm) was loaded on the IFC for cell viability assessment. After the cell loading step, the capturing efficacy was measured using a Nikon automated microscope.

7 µL of lysis/ATAC mix (0.15% NP40, 1.5xTagment DNA Buffer, 1.5xNextera®

Tagment DNA enzyme 1 (Nextera DNA library preparation kit, Illumina), 1.5xC1 No Salt Loading Reagent (Fluidigm), 1.5U RNasin@ Plus RNase Inhibitor), 7 µL of inactivation mix (8.75mM dNTP mix, 8.5µM Bio-C1-P2-T31 primer (Supplementary table 5), 1xC1 No Salt Loading Reagent (Fluidigm), 2.6U RNasin@ Plus RNase Inhibitor, 41.25mM DTT, 18.75mM EDTA), 7 µL of reverse transcription mix (2.93x SSIV Buffer (Invitrogen), 1xC1 No Salt Loading Reagent, 3.5U RNasin@ Plus RNase Inhibitor, 35U SuperScript™IV Reverse Transcriptase (Invitrogen), 7.995µM C1-P2-RNA-Tso primer (Supplementary table 5), 22.05mM $MgCl_2$) and 24 µL of cDNA-PCR mix (1.233xNEBNext® Ultra™ II Q5 PCR master mix, 1.042µM Bio-C1-P2-PCR-2 primer (Supplementary table 5), 1xC1 No Salt Loading Reagent) were loaded onto the designated wells of IFC, according to the 'ASTAR: ASTAR (1861x/1862x/1863x)' script. The script takes approximately 6 hours. On the IFC, the lysis and Tn5 transposition reaction was first performed at 37°C for 30 min, followed by inactivation of Tn5 by EDTA at 50 °C for 30 min and priming of poly T primers for reverse transcription at 72 °C for 3 min, 4°C for 10 min, and 25°C for 1 min. Next, reverse transcription was carried out at 50°C for 60 min, followed by the heat inactivation of SSIV reverse transcriptase at 80°C for 10 min. In the reverse transcription mix, $MgCl_2$ was added to quench excess EDTA from the previous inactivation step. Then the double-stranded cDNA was amplified using the following conditions: 98°C for 3 min; and 5 cycles at 98°C for 20s, 58°C for 4min, and 68°C for 6min, with a final extension step at 72°C for 10 min. In cDNA-PCR mix (Bio-C1-P2-PCR-2). The use of biotinylated primer for cDNA amplification ensured the cDNA were labelled with biotin, which enabled the segregation of cDNA from the ATAC-DNA by pulldown using streptavidin beads. The ATAC-DNA and amplified cDNA were harvested in a total of 5.5µL of C1 Harvest Reagent (Fluidigm).

### 2. off-IFC procedures of ASTAR-seq

#### 2.1 Separation of ATAC-DNA and cDNA

Samples were transferred from the IFC to a 96-well PCR plate (Harvest plate) after running the 'ASTAR: ASTAR (1861x/1862x/1863x)' script. First, 1mL of streptavidin magnetic beads (Dynabeads™ MyOne™ Streptavidin C1, Invitrogen) were washed twice and resuspended in 550µL of 2X Binding and Washing buffer (2M NaCl, 10mM Tris-HCl (pH 7.5), 0.02% Tween-20). Then, 5.5µL of washed beads were mixed with 5.5µL sample in 96-well plate and incubated at room temperature for 20 min on a rotator. 10µL of the cleared supernatant (ATAC-DNA) was collected and transferred to a new 96-well PCR plate (ATACseq plate). The beads were washed twice with 100µL of 1X Binding and Washing Buffer (1M NaCl,

5mM Tris-HCl (pH 7.5), 0.02% Tween-20) and once with 1X TE buffer (10mM Tris-HCl (PH 7.5), 0.02% Tween-20). 10 µL of nuclease-free water was added to each well and cDNA was eluted at 90°C for 10min. Eluted cDNA was transferred to a new 96-well PCR plate (cDNA plate 1) for PCR amplification.

**2.2 Further amplification of cDNA and clean-up**

On the cDNA amplification plate (cDNA plate 1), 15µL of cDNA-PCR master mix (1X NEBNext® Ultra™ II Q5 PCR master mix, 1.6 µM C1-P2-PCR2 primer) was added before transferring the cDNA from the Harvest plate. The following PCR conditions were used for amplification: 98°C for 3 min, 9 cycles of 98°C for 20s, 64°C for 30s, and 68°C for 6 min, followed by 11 cycles of 98°C for 30s, 64°C for 30s and 68°C for 7 min, and a final extension at 72°C for 10 min.

AMPure XP magnetic beads (20µL, Beckman Coulter) were added to each well to purify the amplified cDNA, and incubated at room temperature for 10 min. The beads was washed twice with 75% ethanol and dissolved in 10µL of nuclease free water. The eluent was transferred to a new 96-well PCR plate (cDNA plate 2)

**2.3 Preparation of ASTAR RNAseq libraries for Next Generation Sequencing**

To quantify the average cDNA concentration in the size range of 200-9000 bp, cDNA of 11 cells from cDNA plate 2 were run on an Agilent High Sensitivity DNA Chip. cDNA was further diluted to a final concentration of 0.15-0.2 ng/ µL for mRNAseq library preparation following "Using C1 to Generate Single-Cell cDNA Libraries for mRNA Sequencing" manual. Briefly, 1.25 µL of diluted cDNA was added to 2.5µL Tagment DNA buffer and 1.25µL Amplification Tagment Mix (Nextera XT DNA Library Prep Kit) and incubated at 55°C for 10 min for tagmentation, followed by neutralization with 1.25µL NT Buffer. 3.75µL NPM and 1.25µL of two Nextera® XT indexes were added to each well for barcoding cDNA from each cell. The PCR program was as follows: 72°C for 5min, 95°C for 30s, 22 cycles of 95°C for 10s, 55°C for 30s and 72°C for 1min, and a final extension at 72°C for 5 min. All PCR products were pooled together at a final volume of ~ 1.1 mL. The pooled ASTAR RNAseq libraries was purified twice with AMPure XP magnetic beads (Beckman Coulter, 0.75X). Quality control of the ASTAR RNAseq libraries was assessed by both Agilent bioanalyzer result and qPCR on specific genes.

**2.4 Preparation of ASTAR ATACseq libraries for Next Generation Sequencing**

10µL of supernatant was added on to ATACseq plate with 90µL of ATAC-PCR master mix (1X NEBNext® Ultra™ II Q5 PCR master mix, 1.14 μM custom Nextera dual-index PCR primers (Supplementary table 5)). The PCR program was as follows: 72°C for 5min; 98°C for 30 s; and 22 cycles of 98°C for 10 s, 72°C for 90 s and a final extension at 72°C for 10 min. The PCR products were pooled at a final volume of ~ 9mL. ASTAR ATACseq libraries were precipitated by mixing with 32.2 mL of 100% Ethanol and 4.6 mL of 3M sodium acetate, and incubating at -80°C for overnight. The mixture was centrifuged at 15000g for 20 min at 4°C, and the pellet was washed with 75% ethanol twice and resuspended in 200μl nuclease free $H_2O$. The ASTAR ATACseq library was purified by MinElute column purification kit and eluted in 50 μl Nuclease-Free $H_2O$. Next, the ATACseq library was purified using AMPure XP magnetic beads (Beckman Coulter). ASTAR ATACseq library was incubated with AMPure XP magnetic beads (1.2X) at room temperature for 10 min, washed twice with 75% ethanol, and eluted in 30 μL Nuclease-Free $H_2O$. Quality of the ASTAR ATACseq library was assessed by Agilent bioanalyzer.

**ASTAR-seq Library Sequencing**

192 ASTAR-mRNAseq libraries were sequenced on a lane of HiSeq 4000 sequencer by 101 bp pair-end sequencing. 192 ASTAR-ATACseq libraries were sequenced on a lane of HiSeq 4000 sequencer by 50 bp pair-end sequencing.

### BIOINFORMATIC ANALYSIS

#### RNA-seq Analysis:

**Mapping of scRNA-seq Libraries:**

The scRNA-seq libraries were mapped to mm9 (for mouse libraries) and hg 19 (for human libraries) using star aligner[31]. We allowed up to 2 mismatches and removed reads that map to more than one locus. The option "–outSAMstrandFiled intronMotif" was used to make the bam outputs of star compatible with subsequent analyses. Detailed script is shown in Supplementary File 1 - Bioinformatic scripts.

**Filtering of scRNA-seq Libraries:**

The bam outputs were uploaded to SeqMonk (https://www.bioinformatics.babraham.ac.uk/projects/seqmonk/). The RNA-seq QC plot was generated. For mouse samples, libraries with gene detection rate below 15%

and/or exon mapping percentage below 75 were filtered out. For human libraries, cells with gene detection rate below 6% were filtered out. If more than one read was detected in an exon of a gene, the gene was counted as a detected gene as per SeqMonk protocol.

**Gene Coverage:**

We merged the bam files belonging to each category of cells using samtools[32] merge. Then we used the geneBody_coverage.py module of RSeQC[33] to determine the distribution of the mapped fragments over the genebodies of house-keeping genes.

**Generation of hg19 and mm9 GTF Files:**

The GTF files were generated using the "genePredToGtf" tool designed by UCSC.

**Quantification and Normalization of the scRNA-seq Libraries:**

The bam files and the generated GTF files were used as inputs for Cuffquant[34]. Options -u and -m were included in the cuffquant script. The abundances files were then used as inputs for Cuffnorm[34]. The classic-fpkm normalization style was used. Detailed scripts are shown in Supplementary File 1 - Bioinformatic scripts.

**Pseudotime Analysis:**

The FPKM table generated by Cuffnorm was used as an input for Monocle[27]. The FPKM values were converted to mRNA counts using the "relative2abs" function[28]. The expression family was set to negbinomial while creating the CellDataSet. Then, Size Factors and Dispersions were estimated with default parameters. ReduceDimension function was used with method= 'DDRTree". This was followed by cell ordering (orderCells) and trajectory plotting. Detailed script is shown in Supplementary File 1 - Bioinformatic scripts.

**Correlation with the Mouse Cell Atlas:**

The FPKM table output of cuffnorm (mouse libraries) was uploaded to Mouse Cell Atlas[17] (http://bis.zju.edu.cn/MCA/blast.html). The output obtained after the MCA anlaysis was used to create PCA graphs using FactoMineR[35] package in R and to categorize the populations of the 2i and mESC ASTAR RNA-seq libraries. The treshold used to consider a cell to belong to a particular lineage was 0.414.

**RCA Analysis:**

The FPKM table output of cuffnorm (human libraries) was used as an input for RCA[36]. The human ASTAR RNA-seq libraries were clustered using the Global panel mode of RCA with default parameters. Detailed script is shown in Supplementary File 1 - Bioinformatic scripts.

**Gene Ontology Analysis:**

Lists of genes of interest were uploaded to DAVID[37] (https://david.ncifcrf.gov/). The terms identified by the "GOTERM_BP_DIRECT" functional annotation were used to generate the bar charts. We ensured that the selected GO terms appear only in one of the tested lists.

**UCSC Genome Browser Screenshots:**

The libraries belonging to each cell type were merged using samtools merge. This was followed by the creation of tag directories using the "makeTagDirectory" script of HOMER[38]. Finally, the "makeUCSCfile" script was used and the options -style and -fragLength were set to rnaseq and given respectively.

**Meta-Analysis (RNA-seq):**

For meta-analysis, FPKM values for Bulk K562 RNA-seq, K562 ASTAR RNAseq, BJ ASTAR RNAseq and BJ scRNA-seq were quantified using SeqMonk (RNA-seq Quantification pipeline). The option "correct for gene length" was chosen. Genes were not converted to log. The values obtained were then used to calculate spearman correlation among the libraries (cor function in R). The correlation values were then subjected to PCA using "FactoMineR" [35] (http://factominer.free.fr/) package in R.

**ATAC-seq Analysis:**

**Mapping of the scATAC-seq Libraries:**

Libraries were mapped to mm9 and hg19 genome using STAR aligner[31] similar to the mapping of the scRNA-seq libraries with the addition of the options --alignIntronMax 1 and --alignEndsType EndToEnd. Detailed script is shown in Supplementary File 1 - Bioinformatic scripts.

**Determination of Highly Accessible Regions (HARs):**

Human and mouse libraries were merged independently using samtools merge[32]. Duplicates were removed using the MarkDuplicates module of PICARD. Peak calling was performed using MACS2[39] and the --nomodel --nolambda --keep-dup all --call-summits options were utilised. The narrowPeaks output of MACS2 was considered as the HARs. Detailed scripts are shown in Supplementary File 1 - Bioinformatic scripts.

**chromVAR Analysis:**

The duplicates were removed from the single-cell libraries using the MarkDuplicates module of PICARD. Reads mapping to chromosome M and Y were removed. These libraries were then uploaded to chromVAR[40] along with the narrowPeaks file as an input for the getCounts function of chromVAR. QC was performed using the filterSamples function. Motif variability over these HARs was measured by using the computeDeviations function followed by the computeVariability option. The scATAC-seq libraries were correlated using the getSampleCorrelation module. t-SNE clustering of the scATAC-seq libraries was carried out using the deviationsTSNE option with a perplexity setting of 30. For meta-analysis, the published scATAC-seq libraries were processed similarly and included in the analysis. Detailed script is shown in Supplementary File 1 - Bioinformatic scripts.

**Further Processing of scATAC-seq Libraries:**

Group information was added to each de-duplicated library using the AddOrReplaceReadGroups module of PICARD. This was followed lexicographical sorting of each library using ReorderSam module of PICARD. The option ALLOW_INCOMPLETE_DICT_CONCORDANCE was set to TRUE. The sequence dictionaries for hg19 and mm9 that were used for sorting lack ChrM and other ambiguous chromosomes. Each library was further indexed using samtools index.

**Nucleosomal Pattern Determination:**

The histograms were generated using CollectInsertSizeMetrics module of PICARD.

**Depth of Coverage Detection:**

Coverage of each processed library over the HARs was quantified using

DepthOfCoverage module GATK[41] tools v3.46. The option COUNT_FRAGMENTS was used. The values indicated in the total_cvg columns of the interval_summary outputs of GATK were used for downstream analysis.

For correlation with published scATAC-seq libraries, the single cells belonging to each study were merged using samtools merge. Then the merged bam files were subjected to further processing and filtering as mentioned above. The coverage values were then used to generate the correlation dotplots. The correlation values were calculated using EXCEL. Detailed script is shown in Supplementary File 1 - Bioinformatic scripts.

**Motif Analysis:**

The findMotifsGenome.pl script of HOMER[38] was executed in order to identify the known motifs enriched in the differentially accessible regions.

**UCSC Genome Browser:**

The single-cell libraries belonging to each individual cell type were merged using samtools merge. Tag directories were created using makeTagDirectory module of HOMER. This was followed by the implementation of makeUCSCfile script[38].

**Bimodal technologies and published scATAC-Seq:**

The scRNA-seq libraries of other bimodal techniques and published scATAC-Seq libraries were mapped using STAR aligner, similar to ASTAR-seq libraries as mentioned above. The log.final.out file of each library was used to determine the depth and mapping percentage. To determine the mapping to exons, the mapped files were uploaded to SeqMonk. The RNA-Seq QC Plot function of SeqMonk was used to identify the mapping to exons percentage in each cell in every technique. Duplication rates were determined using the MarkDuplicate module of PICARD.

**Average Enrichment Profile:**

The average enrichment of the ASTAR ATACseq libraries over tss was determined using ngsplot[42].

**Integrative Analysis:**

CoupleNMF[19] was used to cluster the cells based on the integration of both scATAC-Seq part and scRNA-Seq part. The K setting was determined by running the script starting with k=5 and then determining the best NMF score. We kept on reducing the K value and re-running the script until the NMF score obtained was > 1 (For human K=3 and Mouse K=2). The motifs enriched by the peaks specific to each NMF clusters were determined by findMotifsGenome.pl. For the human libraries, the genes identified to be significantly expressed in each cluster were subjected to cTen[30] (http://www.influenza-x.org/~jshoemaker/cten/) for identifying the lineages they enrich. The mouse libraries were clustered using Seurat[43] to ensure correlation of the ASTAR RNA-Seq with the CoupleNMF clusters. For human libraries, the NMF clusters were superimposed on the pseudotime trajectory. For the mouse ASTAR ATAC-Seq, the cluster-specific regions were used to calculate deviations instead of JASPAR motifs using the "getAnnotations" Function of ChromVAR. The heatmaps of ASTAR ATAC NMF clusters were created by obtaining the reads counts over the NMF cluster-specific peaks using featureCounts[44] with -F SAF option. The values obtained were used to generate the heatmaps using heatmap.2 function of gplots in R.

**Combined tSNE (CoupleNMF)**

The genes and accessible regions that were considered significant by the NMF clustering were subsequently used in a combined matrix to cluster the cells based on both the signals together using Seurat. The accessible regions raw coverage counts were generated using GATK[41], as shown in Supplementary File_Cicero.txt. The expression values of significant genes were added to the same matrix. The matrix was used as input for Seurat to cluster the cells.

**Cis-Regulatory Interactions Prediction:**

ASTAR ATACseq libraries belonging to each NMF cluster were merged using samtools merge to determine the HARs of each cluster as described above. Then coverage of each library belonging to a particular cluster over the HARs of its corresponding cluster was measured using DepthOfCoverage as described above. The raw coverage table was used as an input for CICERO[14]. Cicero Cds were created using make_cicero_cds and then "run_cicero" was performed (with default settings). The plot_connections function was used to visualize the predicted regulatory elements of NMF-identified genes. The coordinates used were 10X zoomed-out from the precise co-ordinates of the gene of interest.

**Network Analysis:**

Genes specific for each mouse NMF clusters were uploaded to STRING[45]. The text-mining option was disabled. The network formed was downloaded in tabular format and then the tables were uploaded to cytoscape[46] for visual formatting. Moreover, each network was subjected to ClusterOne[47] and MCODE clustering[48] to identify the most significant complexes within each overall network. The names of the members of each significant complex were uploaded to metascape[49] to identify the biological process in which this complex is involved.

**Determination of Duplication Rates in ATAC and RNA libraries:**

The mapped files were used as inputs for the MarkDuplicates module of PICARD. The metrics files output of MarkDuplicates contained the information about the duplication percentage. Total reads count and mapped reads count for each library were obtained from the "log.final.out" output of STAR aligner.

**Determination of DNA contamination:**

The fold difference between the enrichment of the merged RNA-Seq and merged ATAC-Seq libraries over the determined HARs was measured using getDifferentialPeaks. The Fold threshold was set to 3 using the –F option.

**Confusion Matrix Generation:**

The confusion matrices were generated using MLSeq[50]. The input for ATAC-Seq was the raw depth of coverage values over HARs. HARs were determined and libraries were de-duplicated as mentioned above (Supplementary File Processing_ATAC_For_ChromVAR.txt). Then GATK version 3.4-46 was used to measure the coverage of each cell over each HAR locus[41], as shown in Supplementary File _Cicero.txt. The input for RNA-Seq was the raw counts table of each gene in each library. The raw counts table was generated using htseq count function[51]. We partitioned the input files randomly into two parts. First part consisted of 70% of the input file and was used as training set. The other part (30% of the data) was used as a test set to determine the accuracy of ASTAR-Seq in distinguishing among various cell types. The training data sets were classified using SVM, Random Forest and PLDA classifiers. The method that demonstrated the highest accuracy was used subsequently to classify the test sets and generate the confusion matrix using the MLSeq functions. Detailed descriptions of the scripts are included as a supplementary file.

**Code Availability:**

Codes for the bioinformatic analysis are available in the Supplementary Information file under ‘Bioinformatic scripts’ folder.
